# Quantifying marine larval dispersal to assess MPA network connectivity and inform future national and transboundary planning efforts

**DOI:** 10.1101/2023.05.01.538971

**Authors:** John Cristiani, Emily M. Rubidge, Patrick L. Thompson, Carrie Robb, Margot Hessing- Lewis, Mary I. O’Connor

**Affiliations:** Fisheries and Oceans Canada, Pacific Biological Station, Nanaimo, British Columbia, Canada, V9T6N7; Department of Zoology and Biodiversity Research Centre, University of British Columbia, Vancouver, British Columbia, Canada, V6T1Z4; Fisheries and Oceans Canada, Institute of Ocean Sciences, Sidney, British Columbia, Canada, V8L4B2; Department of Forest and Conservation Sciences, University of British Columbia, Vancouver, British Columbia, Canada, V6T1Z4; Fisheries and Oceans Canada, Regional Headquarters, Vancouver, British Columbia, Canada, V6C3S4; Hakai Institute, Quadra Island Ecological Observatory, Heriot Bay, British Columbia, Canada, V0P1H0; Institute for the Oceans and Fisheries, University of British Columbia, Vancouver, British Columbia, Canada, V6T1Z4; Santa Fe Institute, Santa Fe, New Mexico, USA, 87501

**Keywords:** ecological connectivity, marine protected area network, larval dispersal, biophysical modeling

## Abstract

A Marine Protected Area (MPA) network, in which multiple reserves are designated in a region, can promote the protection of biodiversity across space. To be effective as a network, the design must consider whether MPAs are likely to be connected through the movement of individuals of species of interest. Additionally, network design may explicitly incorporate design features that promote biodiversity in unprotected habitats through the dispersal or spillover of multiple species. Patterns of dispersal and the ability of MPAs to function as an interacting network, however, are difficult to estimate at broad and transboundary spatial scales, and therefore connectivity is often not fully integrated in the design and assessment of MPA networks. Here, we model the dispersal of multiple nearshore species to estimate the potential connectivity of the existing MPAs in British Columbia, Canada, including connections to MPAs in the United States by simulating dispersal using a biophysical model with regional oceanographic currents. We found that MPAs in BC potentially meet connectivity design criteria for nearshore invertebrate species: the majority of MPAs (65-90%) are likely to exchange individuals (i.e. functional connectivity) and support persistent metapopulations, and more than half the unprotected coast (55-85%) receives a large proportion of the larvae produced in MPAs. Furthermore, we found that species’ dispersal abilities and the level of exposure of an MPA to open ocean can predict dispersal distance when we account for the random effects of dispersal location and season. Therefore, future predictions of connectivity are possible based on these core biological and physical attributes, without running new simulations. Together, these analyses provide a robust and novel assessment of multi-species connectivity that can support the design of new MPAs with transboundary connectivity on the northwest coast of North America.

## 1. Introduction

Ecological connectivity – the exchange of individuals among spatially discrete populations – is a key design principle of marine protected area (MPA) network planning (UN CBD 2010, Carr et al. 2017). Reserves that are functionally connected by the dispersal of marine species promote the persistence of populations and the resilience of biodiversity across space in the face of mounting global pressures (Roberts et al. 2017, Harrison et al. 2020). Despite the recognized importance to effective MPA design, connectivity is often not explicitly considered in the design or assessment of MPA networks because the methods and data requirements for doing so are substantial and often not accessible during the design process (Balbar & Metaxas 2019). As a result, MPA network designs can range from unplanned or opportunistic collections of MPAs without a coherent ecological goal to full networks of interacting MPAs strategically placed to protect various life-stages of multiple species (Grorud-Colvert et al. 2014).

An MPA network is considered *ecologically coherent* when it functions synergistically so that 1) MPAs are connected by dispersal and benefit from each other, and 2) they also interact with and support the wider environment (OSPAR 2006, Jonsson et al. 2020). Implicit within this concept are the source-sink dynamics of metapopulations, where MPAs act as sources of dispersing individuals to other MPAs and to unprotected areas. Metapopulation dynamics can facilitate persistence for populations within and outside the network so that populations protected in MPAs can replenish other populations following disturbance and extreme demographic stochasticity (Kininmonth et al. 2019, Harrison et al. 2020). Although population persistence depends on more than connectivity among MPAs, to meet planning objectives an MPA network should at least minimally contribute to the overall persistence of key species. Quantifying potential connectivity among MPAs can reveal whether a coherent network can emerge from the probable spatial dispersal patterns of marine organisms and it can identify gaps in connectivity for future planning efforts.

MPA networks generally have biodiversity or multi-species goals (Balbar and Metaxas), but the relationship between varying dispersal traits and resulting connectivity patterns is not fully understood (Cowen & Sponaugle 2009, Shanks 2009, D’Aloia et al. 2015), making it difficult to design connected networks for all species (Magris et al. 2016, Melià et al. 2016). In many sessile or low-mobility marine invertebrates and fishes, dispersal occurs during the larval life history stage of development in the pelagic environment, with the duration of time for dispersal referred to as the pelagic larval duration (PLD) (Levin 2006). It would seem intuitive that a longer PLD results in longer dispersal distance (Strathmann et al. 2002, Selkoe & Toonen 2011, Burgess et al. 2016), and “rule-of-thumb” planning approaches based on PLD and dispersal distance have been used to guide the size and spacing of MPAs when more direct measurements of connectivity are not available (Shanks et al. 2003, Gaines et al. 2010, Martone et al. 2021). However, location-specific oceanographic processes (e.g., asymmetric currents), physical characteristics (e.g., wave exposure), and/or behaviour (e.g., vertical migration) can make the realized dispersal distance difficult to predict (Nickols et al. 2015, Daigle et al. 2016, Meyer et al. 2021). For example, in areas with complex coastal topography, hydrodynamics, and variable wave exposure, dispersing organisms may be more likely to encounter a barrier and have their dispersal distance limited (Sponaugle et al. 2002, Adams et al. 2014). Therefore, connectivity estimates to inform MPA design and success are likely to be most effective when they can include local oceanographic models that can capture realized currents and dispersal trajectories and avoid the simple assumption that dispersal period is positively correlated with total distance travelled. A robust assessment of connectivity should go beyond standard spatial configuration and network metrics (e.g. spacing, centrality), and attempt to measure the persistence that can arise from dispersal, as the goal of connectivity as an MPA design principle is to establish ecologically relevant network resilience.

As a signatory of the Convention on Biological Diversity (UN CBD 2010), Canada committed and reached Aichi Target 11 (10% of coast protected by 2020) and has now expanded that commitment to ocean protection by announcing a goal of 30% by 2030 (UN CBD 2021). Connectivity is a stated design principle for Canadian Marine Protected Area Network design (Government of Canada 2011, Government of Canada & Government of British Columbia 2014), but there are currently limited studies that have quantified multispecies connectivity in a robust way for the entire Pacific coast of Canada, particularly for nearshore/coastal systems (see Robinson et al., 2005 and Sunday et al., 2014 for regional studies at different scales). The existing MPAs in British Columbia were implemented on an ad hoc basis and were not designed as a connected network. Currently, MPA network development is being conducted at the scale of large marine bioregions (Rubidge et al. 2016), and within the Northern Shelf Bioregion a development process is underway to integrate new MPAs and turn the existing suite of MPAs into a network with connectivity as a design strategy (MPA Network BC Northern Shelf Initiative 2023). At this broad planning scale, there is an opportunity to evaluate the network effects of dispersal and to incorporate connectivity information in the assessment process as new MPAs are added to the network (Balbar et al. 2020).

Here, we model the potential connectivity of the existing MPAs in British Columbia, Canada by simulating dispersal for multiple nearshore species with typical invertebrate life histories using a biophysical model with regional oceanographic currents. Our analysis of MPA network connectivity consists of three objectives. In the first two objectives, we assess ecological coherence by (1) identifying connected MPAs (i.e., MPA inter connectivity) that potentially create persistent metapopulations, and (2) quantifying the source-population potential of MPAs to unprotected areas of the coast. For each of these objectives, we contrast the results among species dispersal potential (i.e., PLD). We then (3) quantify the influence of PLD and release-site exposure on dispersal distance to determine if connectivity can be predicted with limited biological and physical variables that are expected to influence dispersal. In general, we expect that longer PLDs and increased exposure will correlate with greater dispersal distances but that there will be significant spatial variation to the pattern in this topographically complex region. Together, these three analyses provide a robust, multi-faceted, and high-resolution assessment of multi-species connectivity to support MPA network planning.

## 2. Methods

### 2.1. Study area and MPAs

We focused on the connectivity among coastal MPAs in British Columbia (BC), and we also included the coasts of Washington, Oregon, and southeast Alaska to estimate potential transboundary connections in the greater Pacific Northwest region (Figure 1). The physical and climatic characteristics of this region make predicting connectivity patterns difficult. The coast of BC is topographically complex with many islands and deep inlets (i.e. > 25,000 km coastline with more than 20,000 islands), and seasonal changes to dominant wind patterns and freshwater discharge drive the non-tidal nearshore hydrodynamics on the coast (Thomson 1981).

**Figure 1:**
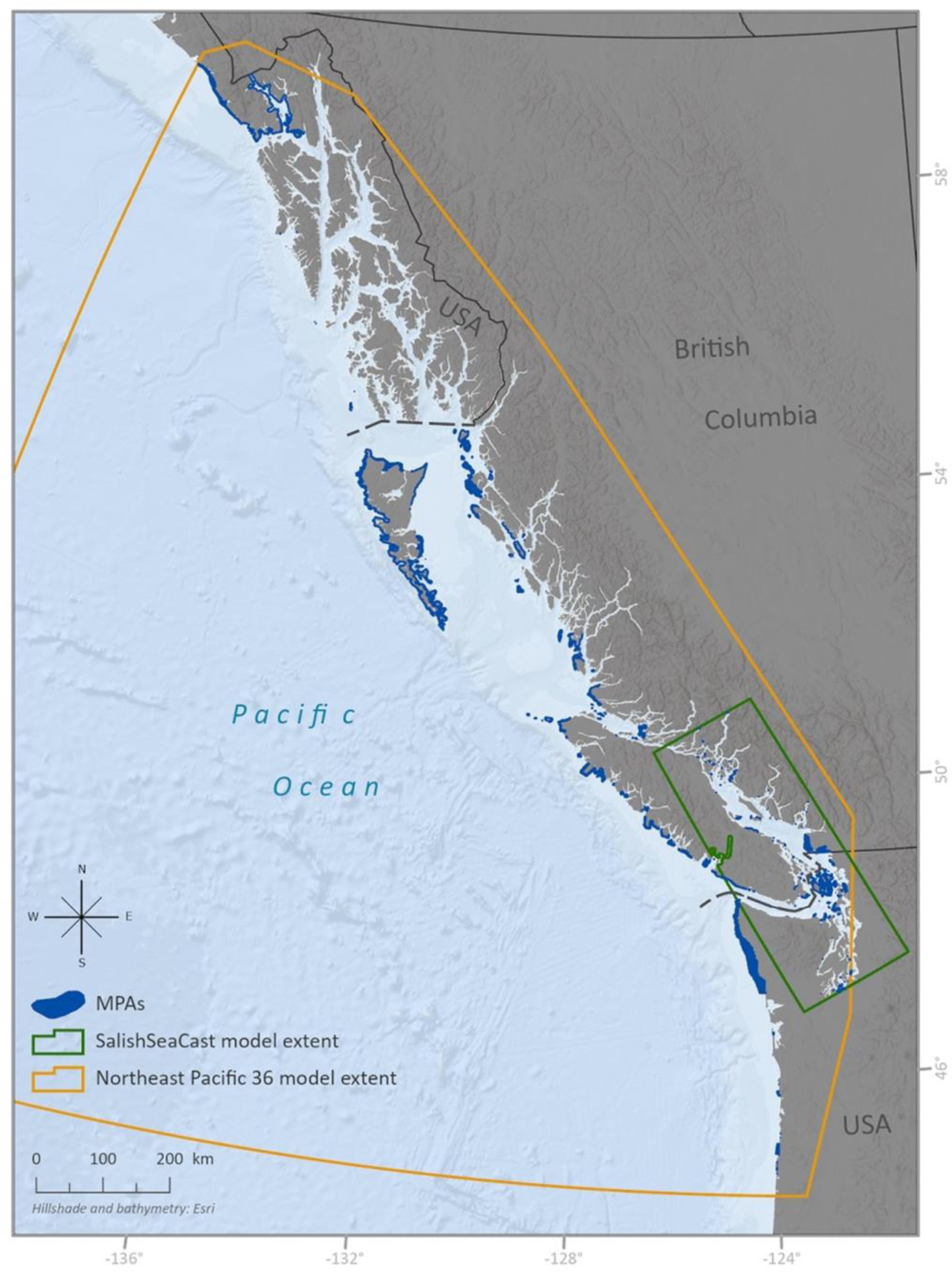
The Pacific Northwest Coast and Marine Protected Areas (MPAs) in British Columbia, Washington, northern Oregon, and southern Alaska. 240 MPAs overlap the study area. Only the nearshore portion of MPAs were included in the analysis (the area of MPAs to a 30-meter depth). The extents of the SalishSeaCast and Northeast Pacific 36 ocean circulation models used to simulate dispersal are shown. MPA polygon size is exaggerated in this map for visualization purposes. Due to the resolution of the Northeast Pacific 36 model resolution, we excluded MPAs from our analysis that are in narrow inlets and fjords where the model is not spatially well resolved.

There are 265 existing MPAs in our study area that meet the IUCN definition of marine protected areas (Day et al. 2019), including Protected Areas, Parks, Conservancies, Ecological Reserves, Wildlife Management Areas, Migratory Bird Sanctuaries, and Other Effective Area-Based Conservation Areas (Canadian Protected and Conserved Areas Database; NOAA MPA Inventory). The MPAs have varying levels of protection and conservation objectives, with some MPAs having general ecosystem objectives, while others may focus conservation efforts on a specific group of species or habitats (e.g., coastal marine bird habitat). Our analysis focused on nearshore species, and therefore we only included the portion of MPAs between a 0-30m depth (limited to the bathymetry fields available in the oceanographic models described below). This eliminated exclusively offshore MPAs (e.g., Hecate Strait Glass Sponge Reefs MPA). We considered spatially discontinuous parts of individual MPAs separately, resulting in 406 spatially distinct MPAs. Lastly, due to the resolution of the oceanographic models (described in the next section), we excluded any MPAs in areas of the models that are not spatially well resolved, such as in narrow inlets and fjords. This resulted in 161 Canadian MPA polygons and 179 USA MPA polygons, for a total of 340 spatially distinct MPA polygons ultimately being included in our analyses (Figure 1).

### 2.2. Biophysical model of dispersal

To simulate dispersal, we built a biophysical model in which species traits (i.e., PLD, dispersal mortality rate) (section 2.2.1.) and oceanographic currents (section 2.2.2.) combine to influence organismal movement (section 2.2.3.) and generate connectivity patterns (section 2.2.4.) (Figure 2). These are potential connectivity patterns that reflect transport and settlement, whereas realized connectivity requires reproducing in a destination location to establish a genetic connection.

**Figure 2:**
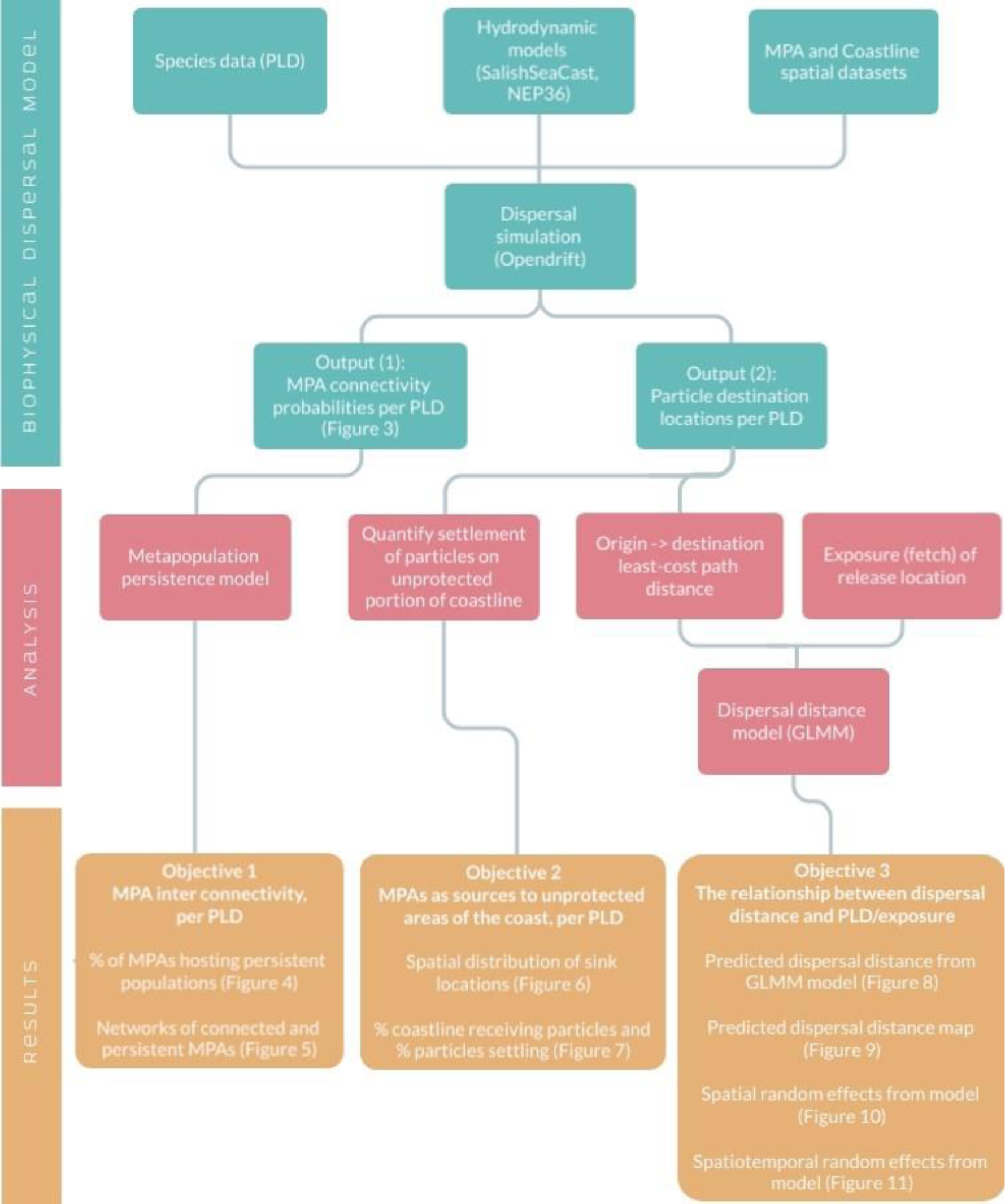
Modeling and analysis approach used to assess dispersal and Marine Protected Area (MPA) connectivity. The biophysical model produces dispersal data for 3 analyses that ultimately produced results on (1) MPA inter connectivity (i.e., MPA to MPA connections), (2) MPAs as sources to unprotected areas of the coast, and (3) the relationship between dispersal distance and Pelagic Larval Duration (PLD) and exposure. For each of these analyses, we contrast the results among species dispersal potential (i.e., eight PLDs).

#### 2.2.1. Focal species – pelagic larval duration

We focused on a selection of nearshore species of commercial, recreational, conservation or cultural importance in BC. We used information on the dispersal drift time of 27 nearshore species from Burt et al. (2014), which included 24 invertebrates (e.g. mussels, scallops, urchins, crabs) and 2 kelp species (e.g. giant kelp and stalked kelp spores). In addition, we included a drift time of 21 days, which is the maximum time that reproductive eelgrass shoots (*Zostera marina*) remain buoyant, allowing for transfer of eelgrass and also enabling rafting of epifaunal invertebrates (Harwell & Orth 2002, Källström et al. 2008, Cristiani et al. 2021). To parameterize our model, these values were binned into eight representative PLD values: 1, 3, 7, 10, 21, 30, 40, 60 days (while our analysis includes species that technically do not have a larval life history phase, for simplicity we used the common term ‘PLD’ to refer to any pelagic drift of a species). We chose not to consider fish species since we are simulating the movement of passive particles with minimal to no swimming ability and our metapopulation model is parameterized for sedentary adult populations (described in the following sections).

#### 2.2.2. Hydrodynamic models

We used two hydrodynamic models to achieve coastwide coverage of adequate resolution: the SalishSeaCast (SSC) and Northeast Pacific 36 (NEP36) ocean circulation models (Figure 1). Both models are configurations of the Nucleus for European Modelling of the Ocean (NEMO) and are forced by wind, rivers, tides, temperature, salinity, precipitation, atmospheric pressure, and radiation fluxes. The SSC model uses an approximately 0.5 km horizontal resolution, covers a smaller spatial extent, and is described in detail by Soontiens et al. (2016), Soontiens and Allen (2017), and Olson et al. (2020). The NEP36 model has an approximately 2.5 km horizontal resolution (Lu et al. 2017) and covers the entire study area. Both models provide velocity fields with a 1-hour temporal resolution. Given the spatial scale of the analysis and computational constraints, only the surface layers (1 m depth) of the models were used, however, because this study focuses on passive dispersal between nearshore areas, we believe that a 2D analysis provides adequate estimates of successful transport between coastal areas. While we acknowledge that a 3D analysis would allow us to measure the influence of larval vertical migration on connectivity, drifting at deeper depths generally acts to retain larvae, and therefore a model with passive surface dispersal is useful for setting the upper bounds of dispersal extent (Selkoe & Toonen 2011).

We did not combine observations where the models overlap but instead used a nested approach in which the higher resolution SSC model is used exclusively for any analysis in its spatial extent. We also made modifications to the coastline layer to allow us to appropriately use these hydrodynamic models for nearshore analyses in otherwise extremely complex and heterogeneous nearshore habitats (i.e. > 25,000 km coastline with more than 20,000 islands). To do so, we simplified the shape of the coastline polygons to match the gridded resolution of the oceanographic models. This then required simplifying the shape of the MPAs to match the modified coastline.

#### 2.2.3. Dispersal simulation

Dispersal was simulated using the Opendrift trajectory modelling software which allows for individual based modeling of dispersal in the ocean by using a Lagrangian particle tracking approach (Dagestad et al. 2018). Using hourly velocity data from the hydrodynamic models, we simulated dispersal for three seasons (winter: Jan-Mar, spring freshet: May-Jul, summer/fall: Aug-Oct) over three years (2011, 2014, 2017). These time periods allowed us to capture variation by year, season, and tidal cycle. Seasonal changes to wind patterns and freshwater discharge may influence connectivity, and by simulating dispersal at different times we also accounted for species that may only spawn or disperse at certain times of the year.

A dispersal simulation was initiated by simultaneously releasing particles from all 340 MPAs. The quantity of particles released from an MPA was assumed to scale with MPA area, and the particle release points were spaced evenly throughout the MPA. We tested the sensitivity of particle fate location to the particle count released following methodology from Simons et al. (2013) to ensure that we released an adequate amount to capture the variation of possible particle destinations in the system. Particles were released every 4 hours for 2 weeks to capture tidal variation. In total 2.7 million particles were released per simulation time period (3 seasons x 3 years).

Particles were tracked as they were advected by velocity fields in the hydrodynamic models. We applied a diffusion rate to represent sub grid-scale turbulence. Rates of 1.5 m^2^ s^-1^ and 20 m^2^ s^-1^ were used for the SSC and NEP36 models respectively. We applied a 15% daily mortality rate by randomly selecting particles to remove from the simulation (Rumrill 1990, White et al. 2014). This rate was assumed to be constant for all PLD groups due to a lack of information on larval mortality during dispersal for most species. Particles that collided with the coastline were considered stranded and stayed in that position for the remainder of the simulation as wave action induced by Stokes drift can increase the probability of stranding (Bosi et al. 2021). Particles were tracked until the end of the PLD being simulated. At that time, any particles that were over an MPA were assumed to settle and make a successful ‘connection’ between the source population and the destination population.

#### 2.2.4. Biophysical model outputs

The three analyses described below relied on two primary outputs from the biophysical dispersal model. (1) The individual particle destination locations were retained for each PLD group if they ended up in the nearshore area. (2) We then calculated a directional probability of connectivity between MPAs (i.e., MPA inter connectivity) by dividing the number of particles that settle on an MPA by the total amount of particles released from the origin MPA. Connectivity probabilities were averaged across all time periods to obtain overall estimates of connectivity per PLD group.

### 2.3. MPA inter connectivity – metapopulation persistence

While a certain level of connectivity may exist between MPAs, it is important to identify if that connectivity generates sufficient immigration to sustain populations when additional population dynamics are also considered (e.g. local population mortality). To achieve our first objective of identifying connected MPAs that potentially form persistent metapopulations, we built a density-dependent stochastic metapopulation model of persistence for each PLD level. In the case of many marine invertebrates, which have dispersing larvae but are sessile as adults, population persistence is achieved through an adequate rate of immigration, so that an adult is replaced by at least one offspring or immigrant over its lifespan. Therefore, we assume that population dynamics are primarily determined by immigration (including local retention) and adult mortality:

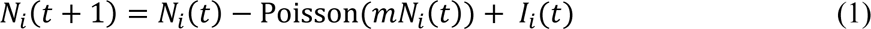

where, *N_i_*(*t*+1) is the population (# individuals) in MPA *i* at time *t*+1, and *N_i_*(*t*) is the population size at time *t*. Mortality rate (*m*) was applied as a constant at each time step (*% t*^-1^) and equal for all populations. To incorporate demographic stochasticity and to ensure an integer population size, we selected values from a Poisson distribution *λ*=*mN_i_*(*t*) at each time point. Immigration (*I_i_*) is the total number of individuals coming into MPA *i* from all other MPAs at time *t*, including MPA *i* (i.e., local retention). The connectivity probabilities from the dispersal simulations (biophysical model sections 2.2.3. and 2.2.4.) were used to determine the probability of an individual arriving from another population:

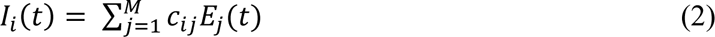

where *M* is the total number of MPAs, and ***c*** is the connectivity matrix among all MPAs. Therefore, c*_ij_* is the connectivity probability from MPA *j* to MPA *i*. Emigration (*E_j_*) is calculated from a simplified constant term (*d*) which combines a reproduction and dispersal rate, such that:

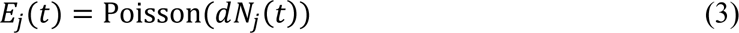

In this model, all the individuals produced through births must disperse but have the chance to recruit back to their home patch. We considered the amount produced and dispersing (*d*) to be viable individuals and not total number of gametes for broadcast spawners, and therefore the rate of individuals dispersing from a patch is analogous to the reproduction rate. To add stochasticity around reproduction and dispersal, and to work with whole individuals, we drew emigration in each time step from a Poisson distribution where *λ*=*dN_j_(t)*. Each individual was assigned to either one of the connected destination MPAs or considered “lost” through a random choice weighted by the modeled connection probabilities (biophysical model sections 3 and 4).

The carrying capacity was set proportional to the area of the MPA, with the largest MPA scaled to have a capacity of 100 million individuals and the smallest MPA to have a capacity of 1000 individuals. These values were chosen so that carrying capacity could be scaled up linearly with MPA area across several orders of magnitude. The values are abstract and do not relate to any specific species’ population. At the end of each time step, population size in each MPA was reduced to carrying capacity (*K_i_*) if the realized *N_i_* exceeded *K_i_*, which was set relative the area of the MPA:

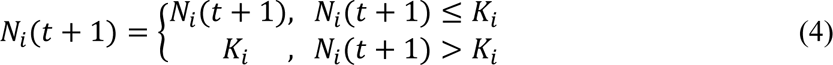

To quantify persistence of each species group (defined by PLD value) in the coastal MPAs, we ran simulations for 1750 timesteps (representing generations), which was sufficient to reach a stable percentage of populations that remained persistent through time. Population sizes (*N_i_*) were initialized at carrying capacity (*K_i_*). To determine the sensitivity of our model to adult mortality (*m*) and dispersal rates (*d*), we ran simulations across 216 scenarios capturing all possible combinations of the 8 PLD values, three levels of mortality rate (0.05, 0.1, 0.15) and nine levels of dispersal rate (0.05, 0.1, 0.15, 0.2, 0.25, 0.3, 0.35). For each scenario, we identified the level of dispersal rate above which the percentage of persistent MPAs saturated. Assuming that there are combinations of mortality and dispersal rates where no MPA populations are persistent or nearly all MPA populations are persistent, saturation of this relationship would indicate a minimum level of likely persistence.

To visualize results, we mapped MPAs to show the spatial scale of persistent metapopulations across PLD values. Using the NetworkX Python package, we calculated the *strongly connected components* formed by the connectivity lines from the dispersal simulations (biophysical model sections 2.2.3. and 2.2.4.) and overlaid these on the map. A component of a network is a cluster of nodes (i.e., MPAs) that are connected to each other but isolated from the rest of the network. A *strongly* connected component of a network is a cluster of nodes where every node is reachable from every other node through directional connections. Therefore, an MPA may be part of the overall component, but if it is only connected as a sink, then it does not connect back to the other MPAs. By examining an MPAs inclusion in a strongly connected component we can see how it contributes to metapopulation persistence or if it only persists as a sink population (Artzy-Randrup & Stone 2010).

### 2.4. MPA source potential to unprotected coastline

To achieve our second objective (Figure 2) of assessing the source potential of all existing MPAs to unprotected areas of the coast per PLD, we quantified (1) the percent of the coastline outside of MPAs that receives dispersing individuals from MPAs, and (2) the percent of total individuals that settle on unprotected coastline. Particles settling on the coastline were rasterized to a 1 km resolution and the sum of particles within each cell was assigned as the cell value. For each PLD level, we calculated the percent of the coastline cells containing particles, as well as the sum of all cells divided by the total amount of particles released. The rasters from the nine time periods were summed to create one overall raster per PLD. Given that one particle settling out of millions released may not indicate significant recruitment from an area, we also ran the calculations for a range of threshold values in which the particles per raster cell required for inclusion in the calculations was increased.

### 2.5. Dispersal distance, PLD and site exposure

To achieve our third objective (Figure 2) of quantifying the influence of PLD and release-site exposure on dispersal distance, we constructed a generalized linear mixed-effects model (GLMM) with spatial and spatiotemporal random fields using the sdmTMB R package (Anderson et al. 2022). We modelled dispersal distance (*Y_s,t_*) for release location *s* and time *t* using a gamma observation model:

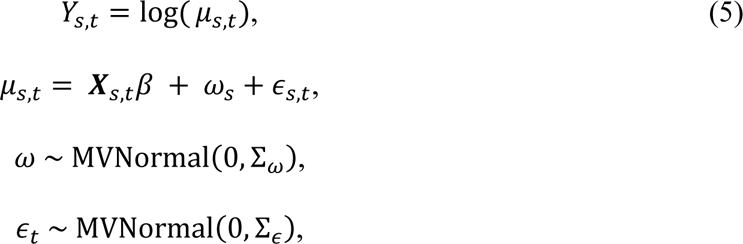

where ***X****_s,t_* is a vector of predictors – PLD and site-exposure (described below), β is a vector of the corresponding fixed effect coefficients, *ω_s_* represents the spatial random field, and *ɛ_s,t_* represents the spatiotemporal random field. These random effects were drawn from a Gaussian random field with covariance matrices Σ_*w*_ and Σ_ɛ_ that were constrained by Matérn covariance functions (Cressie & Wikle 2011). The sdmTMB model constructs a triangulated mesh to approximate a spatial random field with representative locations at the mesh vertices and bilinear interpolation between vertices (Lindgren & Rue 2015). The mesh was constructed with land as a barrier and with a minimum gap of 5 km between vertices to balance predictive accuracy with model fitting runtime. We modeled dispersal distance by season (Jan-Mar, May-Jul, Aug-Oct) and assumed that the spatiotemporal random fields were independent across time (i.e., no temporal autocorrelation between the three seasons modeled). Essentially, the spatial random effects identify spatial patterns that are evident across all seasons, and the spatiotemporal random effects identify patterns that vary from season to season.

To calculate the observed overwater dispersal distance for all particles, based on the origin and destination locations determined from the dispersal simulations, we used a least-cost path analysis with land as a barrier (Figure S1). We only included particles that traveled farther than the resolution of the NEP36 model (2.5 km) since there is higher uncertainty around the exact path of travel within this distance. To quantify exposure, we measured distance to land at 36-degree intervals around a release location and summed the distances (example sites in Figure S2) (Gregr 2014). We expected that sites with low exposure should be more constrained by complex coastal topography (e.g., islands, bay, inlet), and particles from these sites would not travel as far as particles from more exposed sites.

To fit the model, we sampled observed distances for ∼750,000 particles across all PLDs and time periods, and we used an additional 300,000 particles to evaluate the model fit. We compared estimated coefficients for PLD and exposure to infer their relative effect as predictors of dispersal distance. We also compared the spatial and spatiotemporal random effects to assess the variation in dispersal distances that depend on location and time of release and are not associated with our fixed effect predictors.

## 3. Results

### 3.1. MPA inter connectivity

The connection probabilities among MPAs from the biophysical model show the spatial extent of dispersal by PLD (Figure 3). These values act as input to the metapopulation model where we assess how network connectivity and persistence emerges from these individual connections (objective 1). We found that 65-90% of MPAs host persistent populations (Figure 4), with the precise estimate depending on PLD and with shorter PLDs associated with higher persistence rates. Because not all dispersing larvae connect to MPAs, the percent of persistent populations did not saturate until dispersal exceeded mortality (*d* > *m*) by more than 10%. For shorter dispersal periods, fewer particles were lost because they were more likely to settle at home or less likely to drift past nearby MPAs before settling. Spatially, the clustering of persistent MPAs is relatively restricted to adjacent MPAs for a PLD of 1 day, whereas for a PLD of 60 days, nearly the entire coast of BC is potentially connected as an interacting network (Figure 5). Regarding transborder connectivity, for short drift periods (e.g., kelp spores) MPAs in the Salish Sea are more connected to US MPAs than to the central coast of British Columbia, but MPAs on the north coast are only connected to Alaskan MPAs for the highest PLDs (e.g., sea urchin) (Figures 3 and 5).

**Figure 3:**
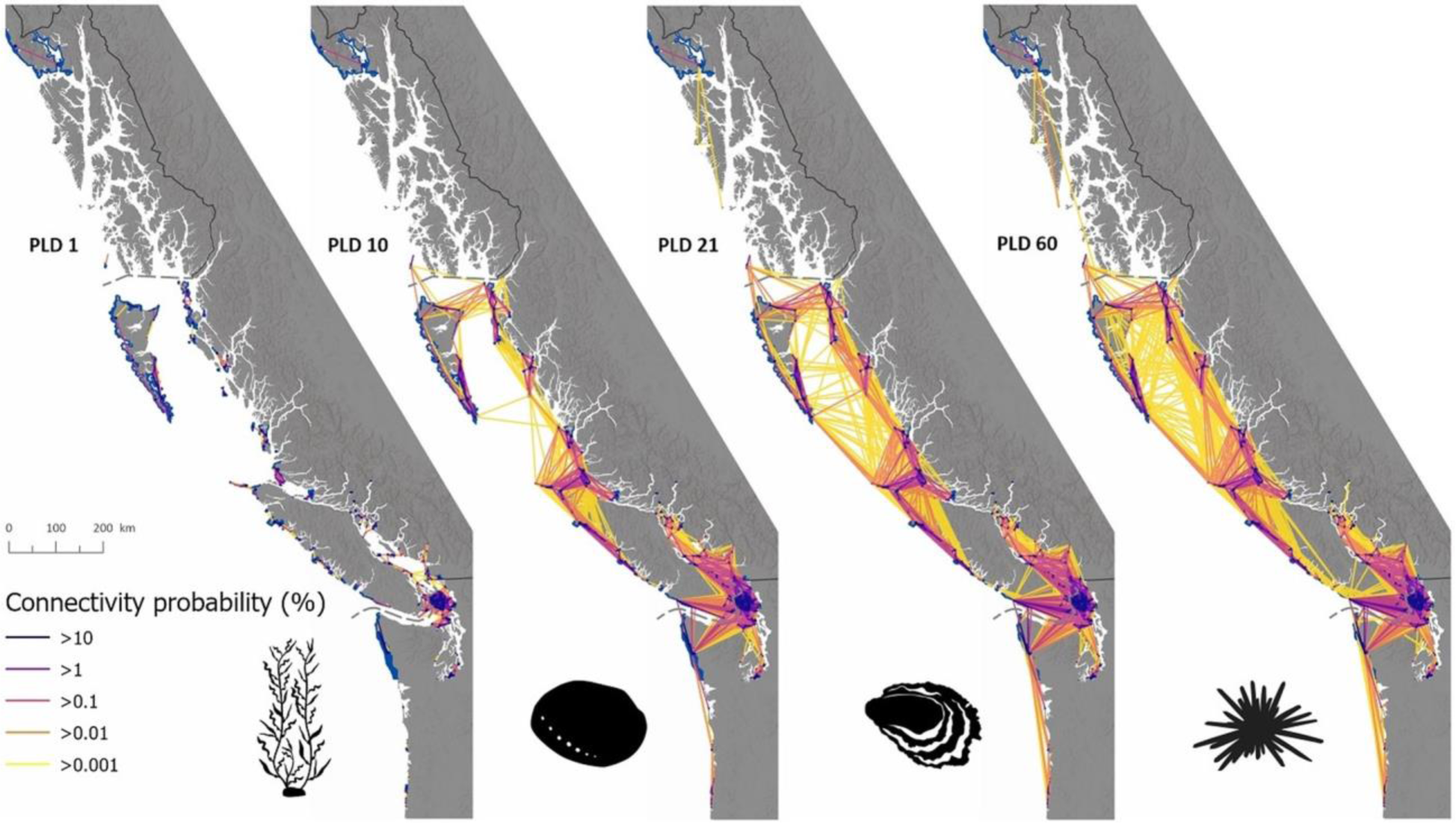
Marine Protected Area (MPA) to MPA connection probabilities, as derived from the particle fates in the biophysical model. Selected species are shown to represent PLD groups. Only four of the eight tested PLDs are shown. Other PLD groups are visually similar. Connectivity results for each PLD are averaged across the yearly and seasonal time periods to represent an overall average. (Open-source image credits: giant kelp – Jane Thomas, abalone – Taro Maeda, Pacific oyster – Natasha Sinegina, sea urchin – Jake Warner).

**Figure 4:**
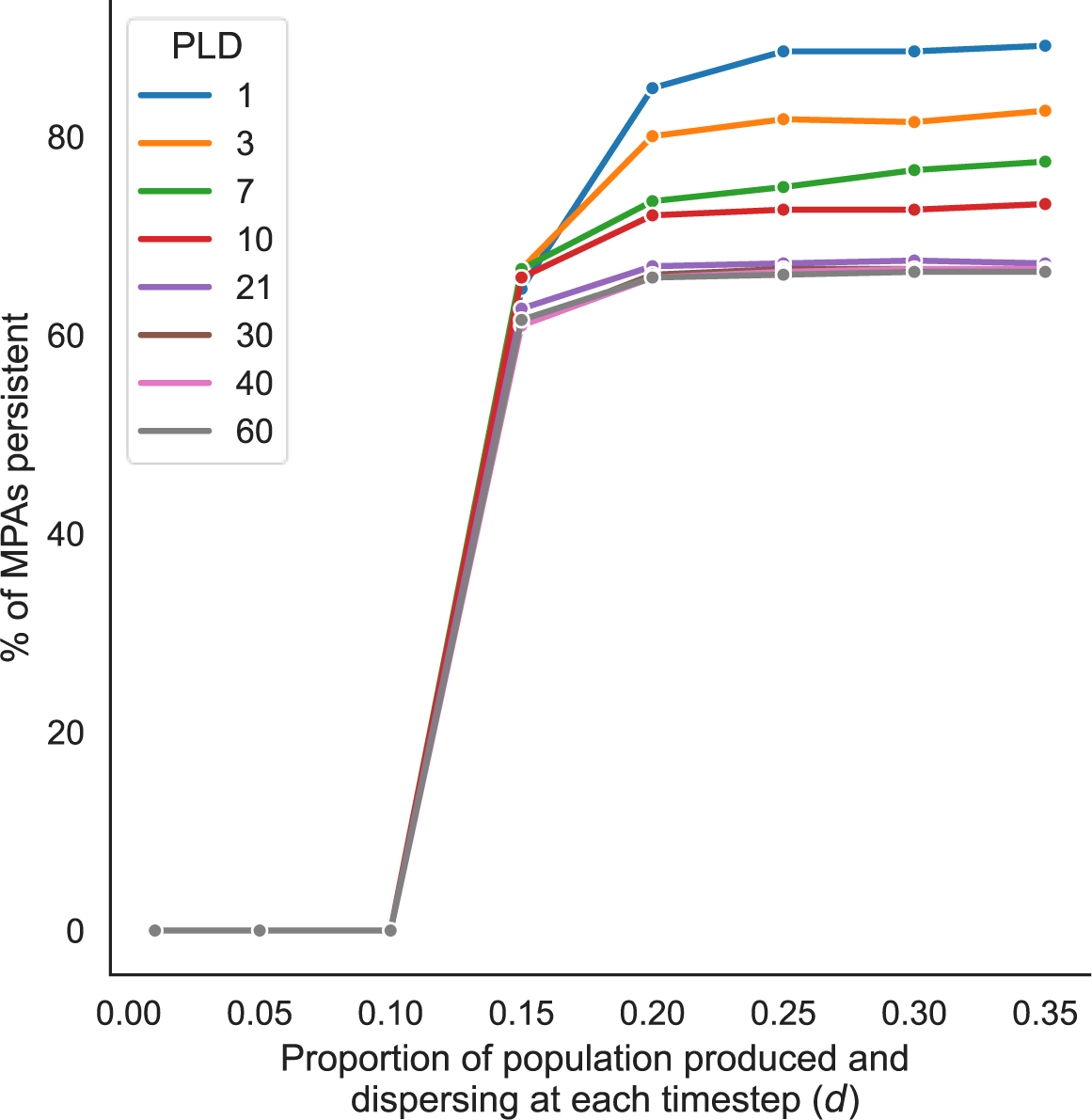
The percentage of MPAs that potentially host persistent populations vs the proportion of the population that is produced and disperses at each timestep (d), based on our metapopulation model. In the case of marine invertebrates with pelagic larvae, all the individuals produced must disperse (but have the chance to recruit back to their home patch). We simplified the amount produced (d) to be viable larvae and not total number of gametes for broadcast spawners, and therefore dispersal rate is analogous to reproduction rate. The results shown are for an adult population mortality rate of 10%, and the stabilization percentage is after 1750 timesteps. For all combinations of dispersal and mortality rates we tested, a stable plateau eventually emerges for all PLD levels once dispersal rate exceeds mortality rate. While a certain percentage of viable larvae disperse at each time step, they will not all connect to MPAs. Therefore, saturation of the % persistent is not achieved until the percent dispersing sufficiently exceeds mortality by > 10%.

**Figure 5:**
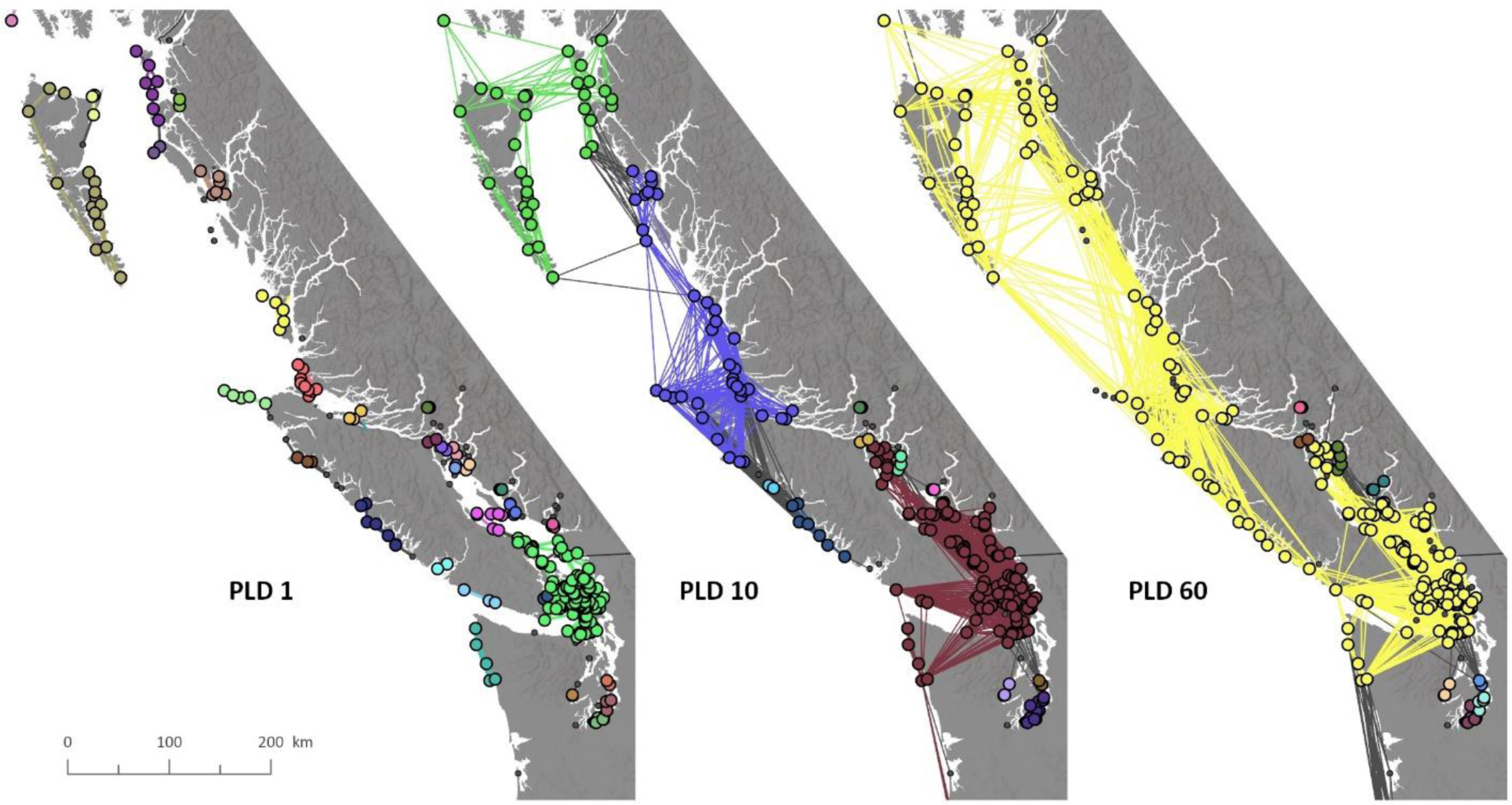
Persistent MPAs as identified from the metapopulation model for three PLD values that represent the range in persistence and extent of components. Points represent the centroid of the MPAs (showing primarily Canadian MPAs for visualization purposes). Separate clusters of MPAs are coloured by their inclusion in a strongly connected component, in which each MPA can be reached from every other MPA. Colors between maps are not related. An MPA can also be persistent through larval retention or as a sink to a component but does not contribute to that component. Small grey points are not persistent.

### 3.2. MPA source potential to unprotected coastline

To quantify the source-population potential of the existing MPAs to unprotected areas of the coast (objective 2), we summed the particles from the biophysical model that recruit to nearshore areas of the coast (Figure 6). We found that 55-85% of the coastline received individuals that disperse from MPAs, depending on PLD (Figure 7a). This percentage is based on at least 1 individual settling in a 1 km^2^ area. If the number of individuals required to create an ecologically significant connection is increased, however, the percent of the coast covered drops quickly. We also found that 25-33% of the total particles released from MPAs recruit to unprotected nearshore areas (Figure 7b).

**Figure 6:**
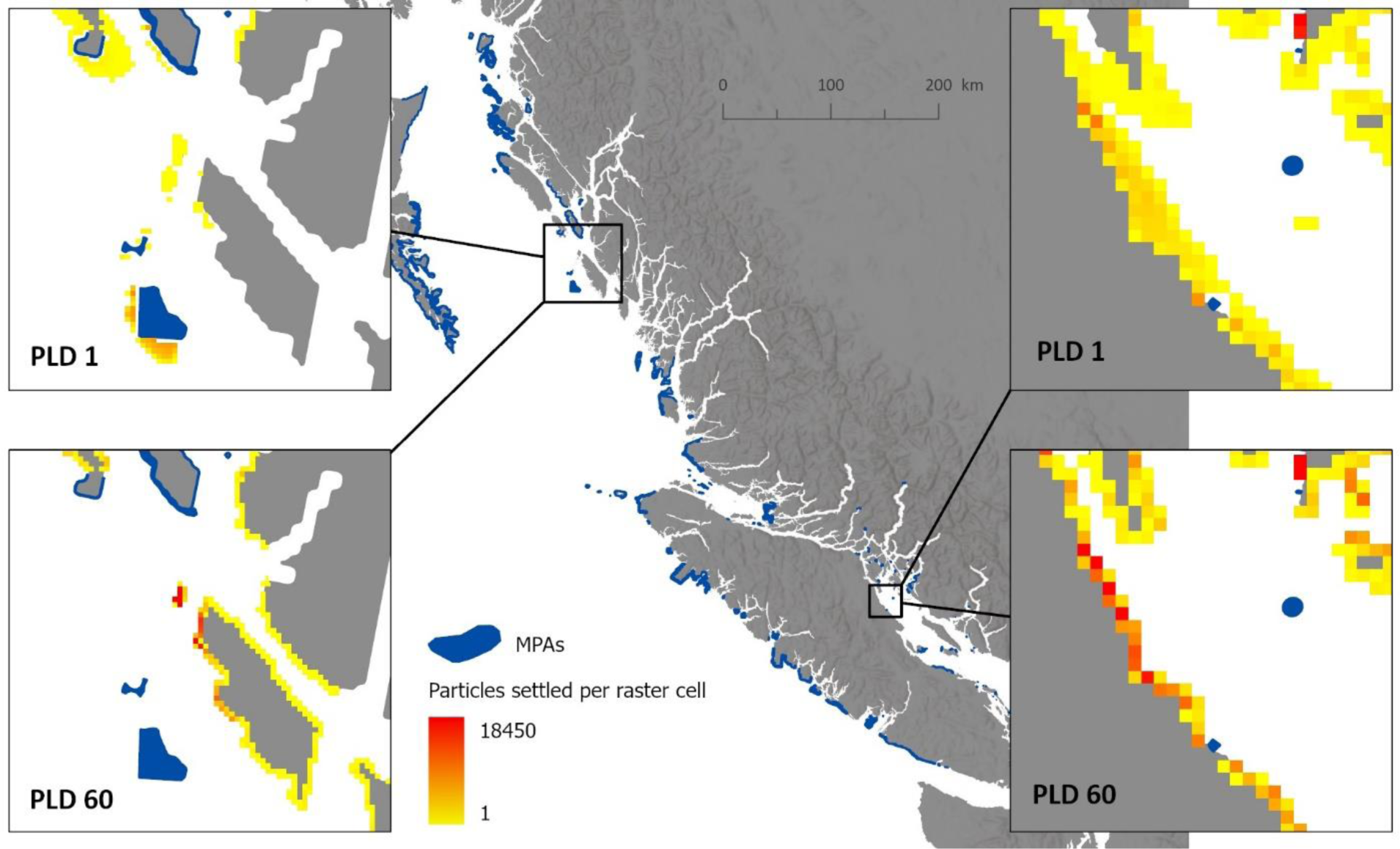
The quantities of particles that recruit from MPAs to unprotected nearshore areas of the coast.The quantities are gridded to a 1 km2 area, and we show results for pelagic larval durations (PLD) of 1 and 60 days to demonstrate the range of values. Inset maps are shown for example because results are not visually evident at the coast-wide scale; the locations featured do not imply any significance. The analysis was done only for the Canadian coastline, and the percentage of coast that this analysis is based on does not include fjords on the central and northern coast where the hydrodynamic model is not sufficiently resolved.

**Figure 7:**
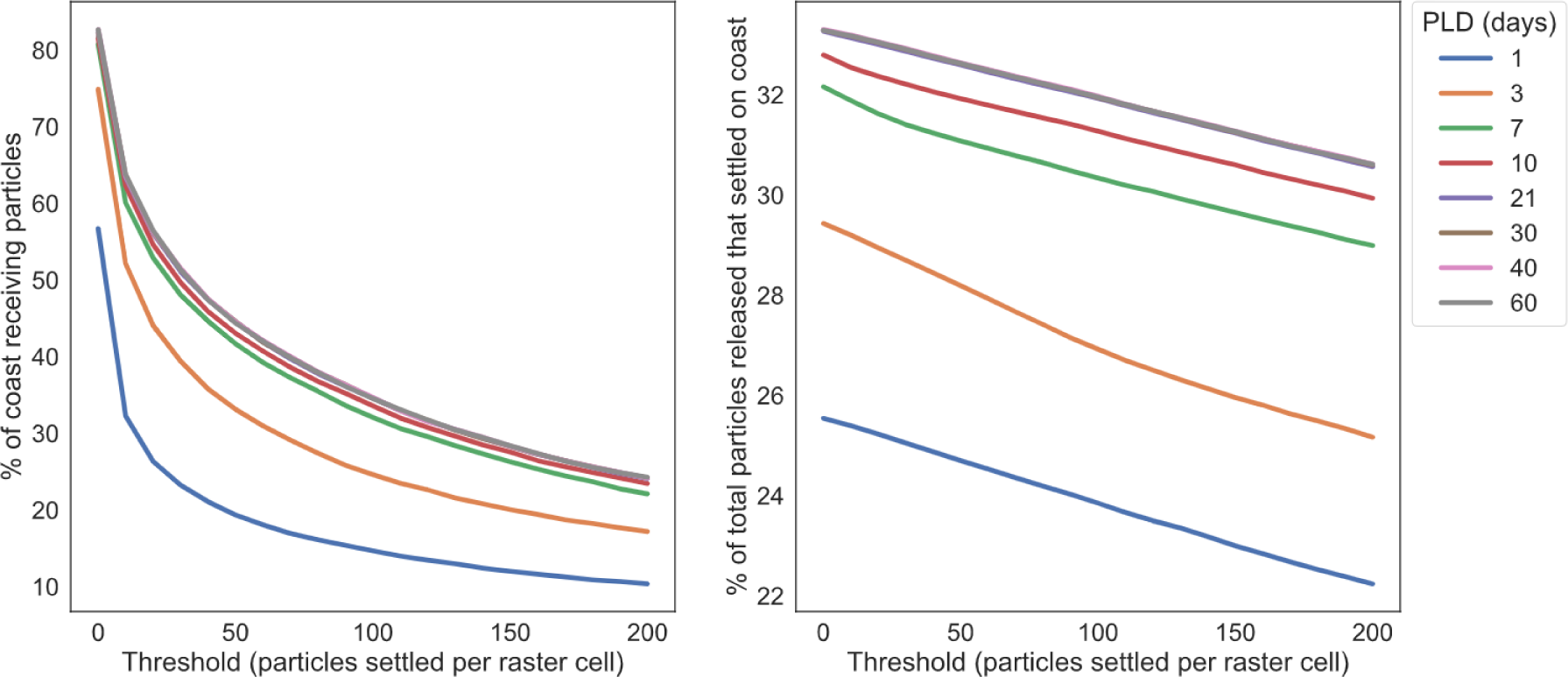
The percentage of the unprotected coastline that receives particles from MPAs, and the percentage of total particles released from MPAs that settle on the unprotected portion of the coast.

### 3.3. Dispersal distance from MPAs

Our GLMM was able to predict dispersal distance from MPAs in data not included in the model with a predictive accuracy R^2^ of 0.35. We found that PLD and exposure of release site explain patterns in dispersal distance from MPAs (objective 3). There was a statistically significant (confidence intervals not overlapping zero) and positive relationship between dispersal distance and the predictors in our spatiotemporal GLMM (Table 1). Thus, the highest dispersal distances occurred for larvae with the highest PLD (60 days) leaving from areas with high exposure (Figure 8). While the relationship between the fixed effects and dispersal distance was statistically significant, there was still considerable variation in the relationship (Figure S3) with the random effects explaining 94% of the variance (Table S1). Therefore, our model is well-suited for making predictions for the BC coast, but less suited to make predictions outside the study area. The spatial random effects indicate that there is additional variation in dispersal distances that depend on geographic location of the origin MPA (Table 1). For example, at the southern end of Vancouver Island, larvae travel systematically farther than predicted by PLD and exposure (Figure 9 and 10). While the release site exposure is moderately constrained, directional currents in this region can transport larvae to the outer coast without encountering coastline in the Strait of Juan de Fuca. Furthermore, the spatiotemporal random fields indicate that there is also geographic influence that varies depending on the month in which the larvae are released. For example, there are notable locations such as the west and east coast of Vancouver Island that switch direction in the magnitude of effects between the winter and summer months (Figure 11).

**Figure 8:**
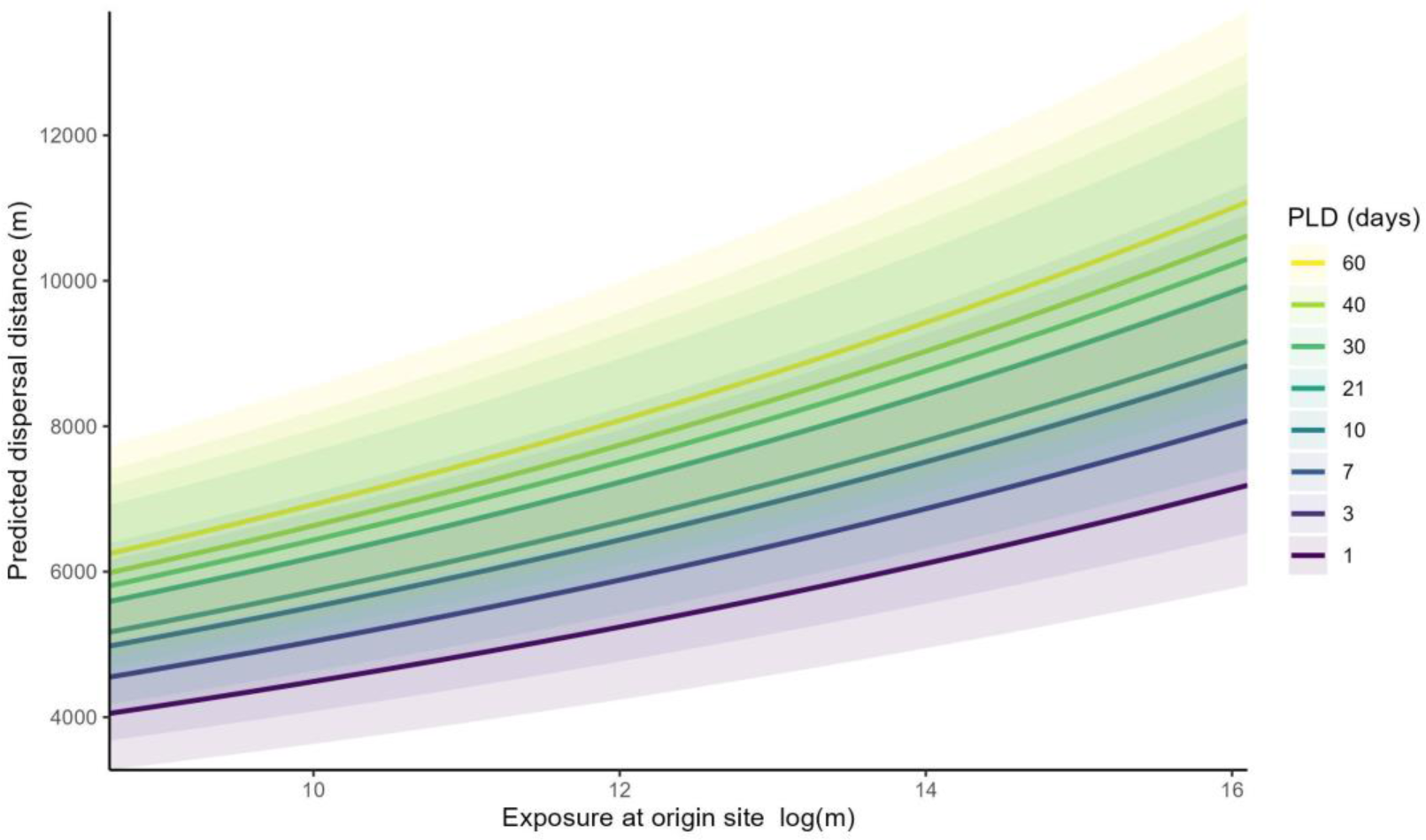
The predicted distance a particle will disperse to other coastal areas based on the exposure at the site of release. Shading indicates a 95% confidence interval. See Figure S2 for explanation of exposure units.

**Figure 9:**
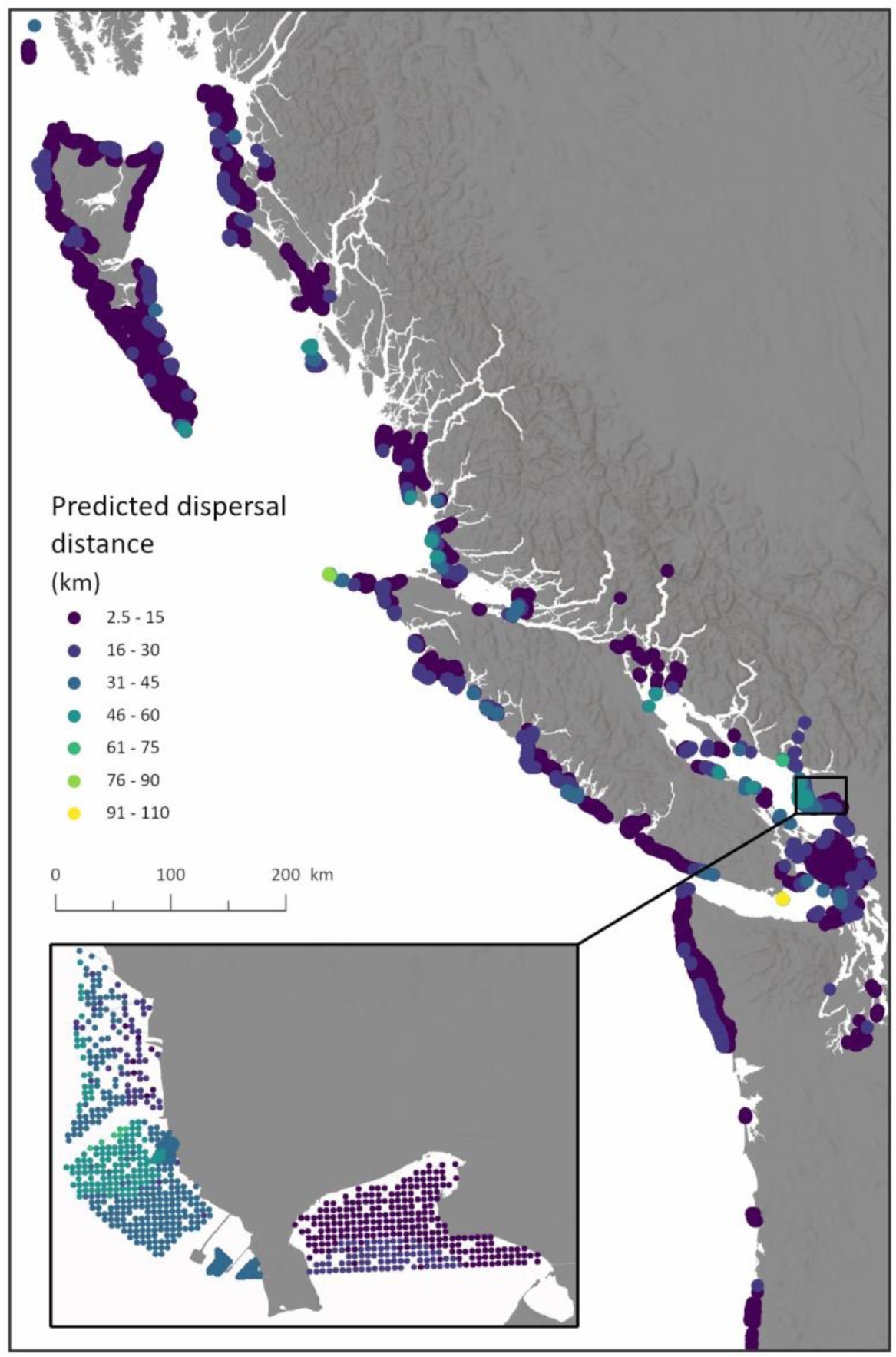
The GLMM predicted dispersal distances to other coastal areas from particle release locations in existing MPAs. The results for PLD 10 are shown to represent an intermediate level of dispersal. Model predictions are based on both the fixed and random effects. Comparing the predicted distances to the influence of spatial and spatiotemporal random effects in Figures 10 and 11 reveal where the fixed effects over/under predict dispersal distance.

**Figure 10:**
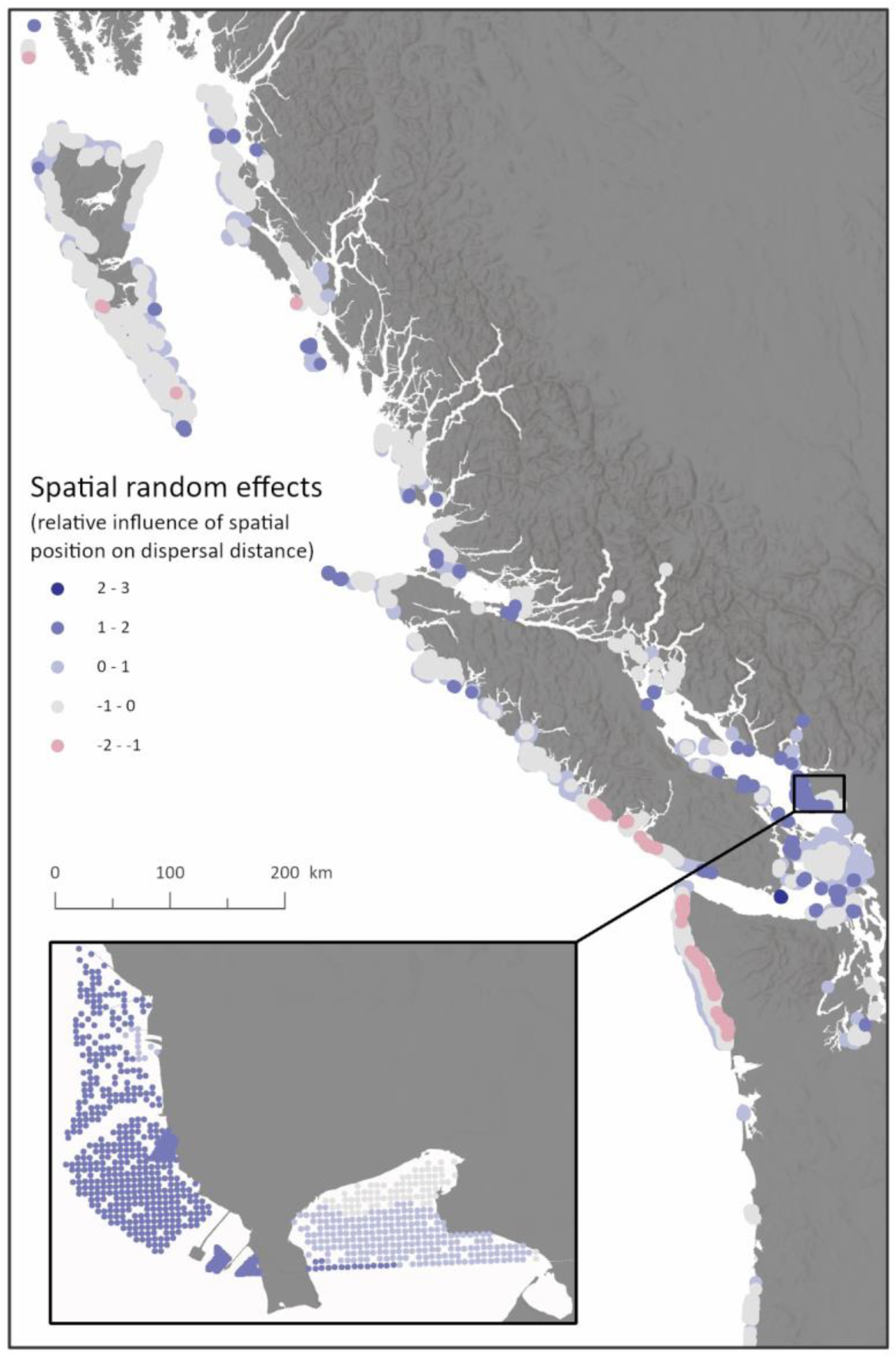
Map of spatial random effects for PLD 10 – relative deviations in space of dispersal distance that are not accounted for by the fixed effects. A positive value indicates that a particle travels farther from that location than can be predicted by just exposure and PLD. A negative value indicates that it travels a shorter distance than predicted by the fixed effects. These deviations represent random spatial variation as well as biotic and abiotic factors not accounted for in the GLMM but that may be affecting dispersal distance.

**Figure 11:**
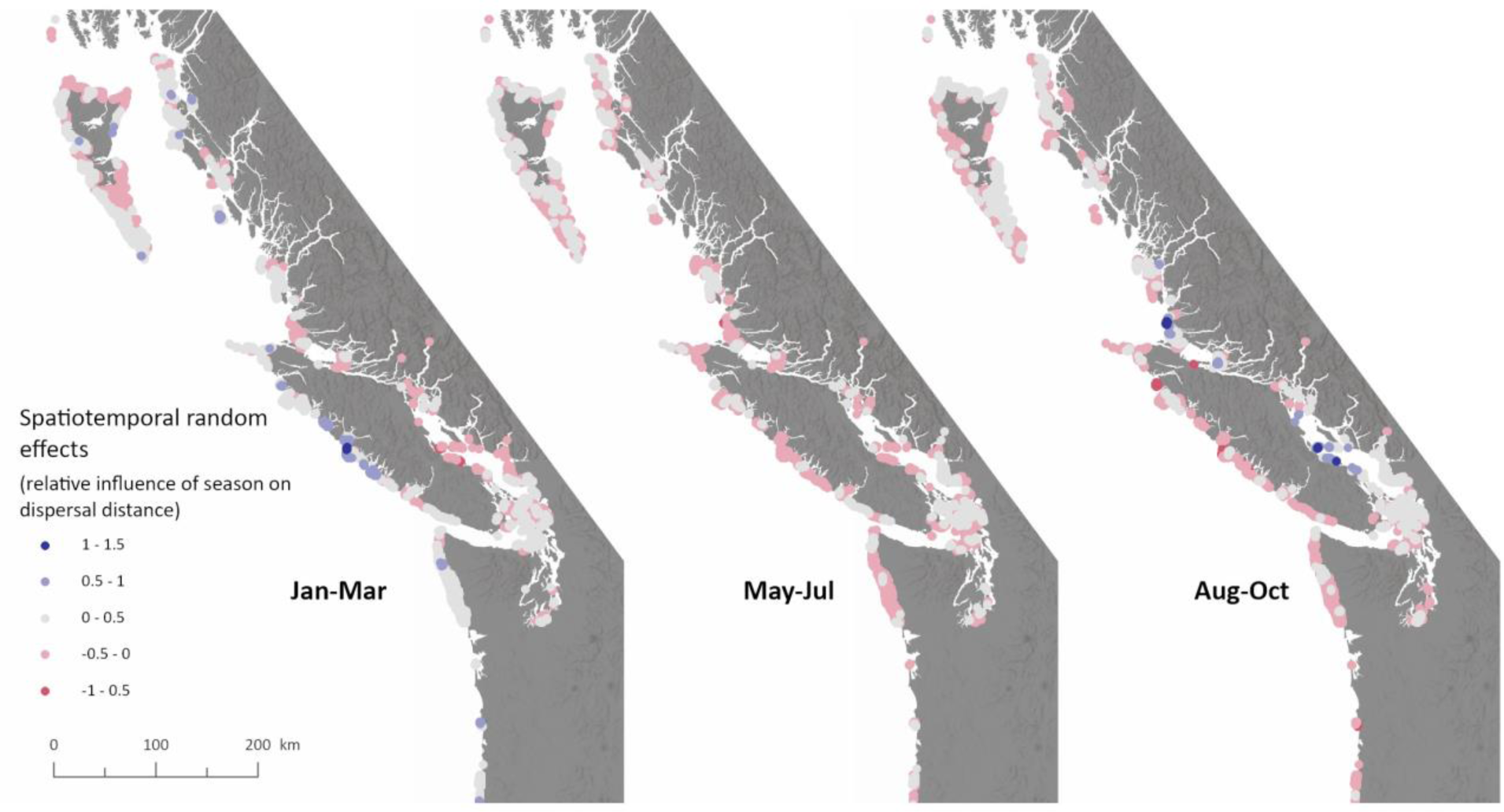
Map of spatiotemporal random effects for PLD 10. These deviations represent random temporal variation as well as biotic and abiotic factors that vary in time and may influence dispersal distance but are not explained by the fixed effects or the spatial random field. Of note is the contrast between January and August, particularly on the west coast of Vancouver Island and the Salish Sea.

**Table 1:** Generalized linear mixed effects model (GLMM) results.The range of the spatial random field is the distance at which two points are effectively uncorrelated in space. Sigma_O and sigma_E are the marginal standard deviations of the spatial and spatiotemporal random fields respectively.

| **Fixed effect terms** | **Estimate** | **CI Lower 95%** | **CI Upper 95%** |
| --- | --- | --- | --- |
| PLD *log(days)* | 0.106 | 0.105 | 0.107 |
| Exposure *log(m)* | 0.077 | 0.071 | 0.084 |
| **Random field parameters** | **Estimate** |  |  |
| Range (km) | 38.362 | 35.203 | 41.804 |
| sigma_O | 6.019 | 5.236 | 6.920 |
| sigma_E | 1.494 | 1.280 | 1.745 |

## 4. Discussion

### 4.1. Evaluating MPA network coherence for multiple species

In this study, we generated patterns of connectivity by simulating dispersal from coastal MPAs to reveal how populations within MPAs potentially interact with each other and the wider environment. We found that the majority of MPAs (65-90%) are likely to exchange individuals and support persistent metapopulations, with PLD having a strong influence on this variation. The majority of the unprotected coast (55-85%) also receives larvae from MPAs. In addition, we found that species’ dispersal abilities and the exposure of an MPA to open ocean can predict dispersal distance when we incorporate the random effects of dispersal location and season. Together, these analyses allowed us to assess the ecological coherence of an MPA network. This approach goes beyond simple metrics of spatial configuration to uncover possible persistence and source-sink dynamics that emerge from reserves being functionally connected, since ultimately, the goal of connectivity as a design principle is to generate network resilience. For example, if an unprotected source to an MPA becomes degraded, it’s crucial that more distant MPAs can provide larval input to stabilize populations. Therefore, it’s necessary to know if an MPA network is self-sustaining, which our model reveals, even if real persistence is enhanced with additional connections from outside of the network (Gaines et al., 2010; Ross et al., 2017). There are currently limited studies showing the benefits of MPA connectivity to population maintenance (Jacquemont et al. 2022), and our study attempts to fill this gap by demonstrating not just that MPAs are connected, but that an ecosystem benefit arises from this connectivity.

We found that the duration of the dispersing period can have a strong influence on the spatial patterns of connectivity in this region. The variation in our results between short and long-distance dispersers emphasizes the importance of considering multiple dispersal abilities in MPA network design. No dispersal strategy connected all MPAs across the study area. While this was not expected over a broad spatial scale and across international borders, it emphasizes that managing a bioregional MPA network requires managing a collection of networks, and the set of MPAs in each network changes with dispersal ability. Longer PLDs achieved lower rates of persistence but had more source potential to other areas of the coast. This may indicate that the spacing of MPAs is not optimal for all dispersal strategies. A species with a long PLD requires more development time in the pelagic environment and may drift past nearby MPAs before it is competent to settle. The longer drift time, however, allowed these species to supply more areas of the unprotected coastline. Therefore, a more consistent spacing of additional MPAs along the coast, and attention to MPA design across coastal boundaries and management jurisdictions, could potentially connect more MPAs for longer distance disperses and also allow shorter dispersers to reach additional unprotected sink habitats.

The recommended spacing for Canadian Pacific MPAs is 20-200 km (Burt et al. 2014, Martone et al. 2021). In an assessment of MPA spacing on the BC coast, Ban et al. (2014) found that most MPAs occurred in clumps, within which MPAs were adequately spaced for expected dispersal distance, but the clumps were separated by distances greater than the recommended spacing. Our results complement these findings and demonstrate the consequences of spatial configuration (e.g., inconsistent spacing) on persistence and source-potential. To study the effects of spacing in more detail, future model iterations can test the range of PLD values that exist for some species (e.g., red sea urchin: 40-60 days), as well as a precompetency trait value (i.e., the *minimum* development time needed to become viable, at which point an organism could settle if it reaches appropriate habitat). This variation in development time can influence dispersal distance and result in varying connectivity patterns for a species (Cecino & Treml 2021). However, since we tested a range of 8 PLD values between 1 and 60 days, we feel we have indirectly captured this variation.

The spatial extent of persistent networks is significantly different between short- and long-range dispersers. While our metapopulation persistence model makes several simplifying assumptions, it demonstrates how dispersal influences persistence, and the resulting patterns match known fundamental network characteristics that generate persistence. Essentially, persistence depends on the connectivity between MPAs forming loops so that a supply of larvae “return” to the origin population from multi-generational and multi-patch movement (Hastings & Botsford 2006, Artzy-Randrup & Stone 2010, Dedrick et al. 2021). An isolated population can therefore persist only if larval retention and survival exceed adult mortality (Burgess et al. 2014). While reproduction and mortality rates influence the level of stable persistence, for species in which larvae are pelagic and disperse, potential persistence of the population can be explained by essential topological characteristics of the dispersal connections: to persist a population must either: (1) be part of a loop (i.e., a strongly connected graph component), (2) a sink of a strongly connected component, or (3) have high local retention. Given that local retention is crucial for isolated populations, future planning could focus on protecting additional sources of larval supply for these populations to increase redundancy in the system.

### 4.2. Predicting dispersal distance from MPAs to aid future design

We constructed a simple model to predict dispersal distance that incorporates one biological characteristic (PLD) and one physical characteristic (release site exposure). While we found a relationship between dispersal distance and these two predictors, there was significant variation attributed to spatial and spatiotemporal random effects. This suggests that there are potentially abiotic factors varying in space and time that are not explicitly included in our statistical model but that are captured by our spatial and spatiotemporal random effects. The unexplained variation attributed to spatial and spatiotemporal random effects is unsurprising given that the relationship between exposure and asymmetric currents is complex, and therefore a low exposure site could also have strong currents that direct movement to open ocean more than predicted by exposure alone (Sponaugle et al. 2002, Adams et al. 2014, Nickols et al. 2015). Numerous oceanographic drifter and simulation studies in the region have found significant spatial and temporal differences in coastal hydrodynamics due to seasonal variation in wind and river discharge (Masson & Fine 2012, Halverson & Pawlowicz 2016, Pawlowicz et al. 2019, Blanken et al. 2020). These seasonal differences have been shown to alter predicted connectivity patterns in BC (Xuereb et al. 2018, Cristiani et al. 2021). These findings emphasize the complexity of modeling and predicting dispersal distance on a topographically complex coastline with spatiotemporally varying ocean currents. Despite this complexity, we have demonstrated that our model is sufficient to make reasonable predictions of dispersal distance based on PLD, exposure, and with the inclusion of spatiotemporal random fields. Therefore, this model could be used in the design of new MPAs on the BC coast by making initial predictions of connectivity without running additional dispersal simulations, but because the predictive accuracy relies on random fields, the model is not as well suited to make predictions in other regions.

### 4.3. Connectivity and marine spatial planning in British Columbia and beyond

The BC MPA Network Strategy, which details the approach for MPA network planning, includes connectivity in its design principles, which involves protecting source populations and ensuring ecological linkages (Government of Canada & Government of British Columbia 2014). The existing MPAs in BC have been designed at various scales with a range of objectives, and connectivity has only been evaluated *post hoc* or with distance-based proxy measures (Burt et al. 2014, Robb et al. 2015, Martone et al. 2021). Thus, our coastwide study fills this gap by directly linking the process of dispersal to patterns of connectivity.

We have demonstrated that the MPAs in BC appears to meet some network connectivity design criteria for nearshore invertebrate species: the majority of MPAs are functionally connected and support persistent metapopulations, and more than half the unprotected coast receives a large proportion of the larvae produced in MPAs. Future applications of our model can involve determining the placement of new MPAs. This could involve running similar simulations from a candidate location to determine its linkages to the existing network, or to run backwards simulations to identify where a source MPA would need to be placed if it is not practical to protect that specific location. It will also be important to incorporate other life history strategies and predict movements of species with more mobility by incorporating additional parameters (e.g., directional swimming). Lastly, effective MPA network design will need to consider the effects of climate change, as species ranges may shift and no longer align with the configuration of reserves that were designed to protect them (Fredston-Hermann et al. 2018). Forecasted oceanographic models and predictions of physiological changes to larval development can be incorporated into biophysical models to predict these shifts, and an increased focus can be placed on protecting future corridors.

While the focus of this study is on marine spatial planning in Canada, we emphasize the inherent borderless nature of animal dispersal and connectivity. A significant number of connections exist between coastal areas with differing governance (i.e., First Nations traditional territories, provincial/state/federal levels of Canada and the United States). Maintaining these individual marine populations may be crucial to the persistence of the larger metapopulations that span borders, especially for dispersal limited species threatened by climate change (e.g., kelp in the transboundary Salish Sea). Therefore, the success of future ocean planning in the northeast Pacific will rely on co-governance and cooperation along a connected coastline that spans international borders. We demonstrate the importance of considering MPA design at a very large scale (>1000 kms), and we provide the basis for a future transboundary framework of MPA design considerations in this region.

## Data availability

The code and datasets generated for this study are available at: <*a Zenodo archived GitHub repository will be made publicly available upon acceptance of the article*>.

## Acknowledgements

We thank Ben Moore-Maley and Susan Allen for technical support with the SalishSeaCast model, and Li Zhai and Youyu Lu for technical support with the NEP36 model. We also thank Charles Hannah and Katie Gale for project guidance, and Sarah Gergel and Coreen Forbes for feedback on the manuscript.

---

**Figure S1:**
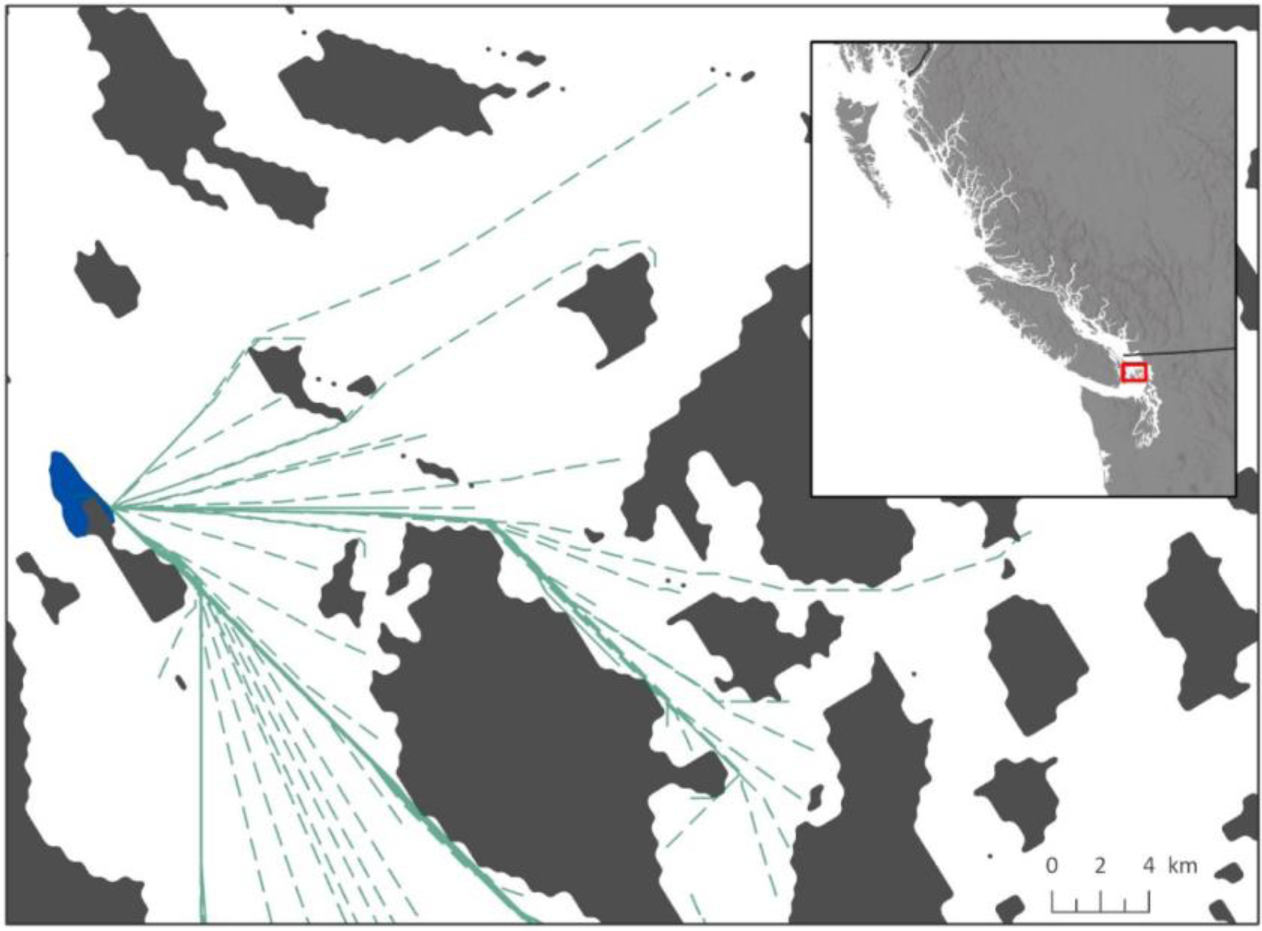
An example of overwater least-cost path distance between a release location in an MPA and the particle destinations.

**Figure S2:**
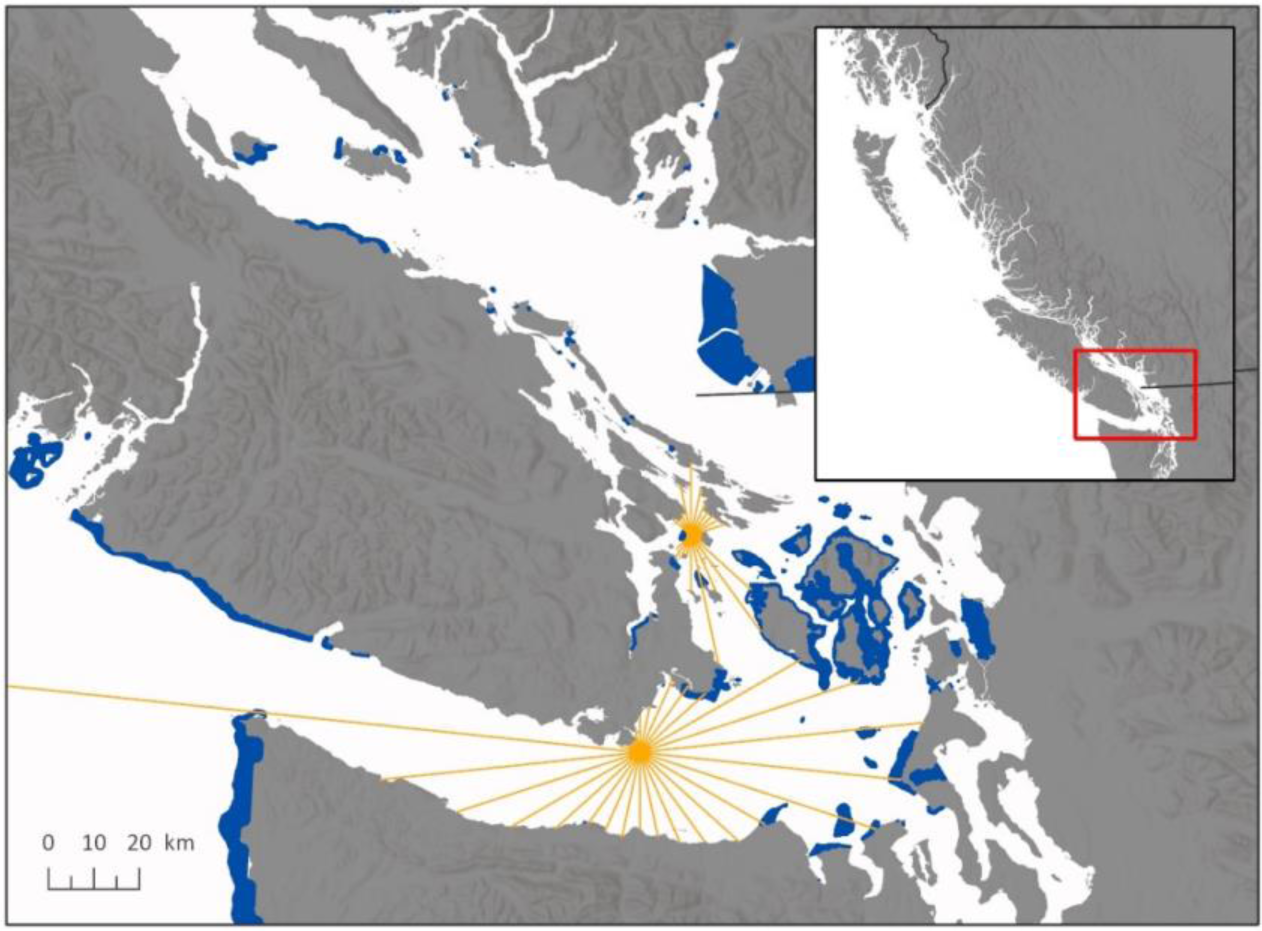
Site exposure lines shown for two release locations. We measured distance to land at 36-degree intervals around a release location and summed the distances. A max distance of 500 km was used to capture any barrier effects.

**Figure S3:**
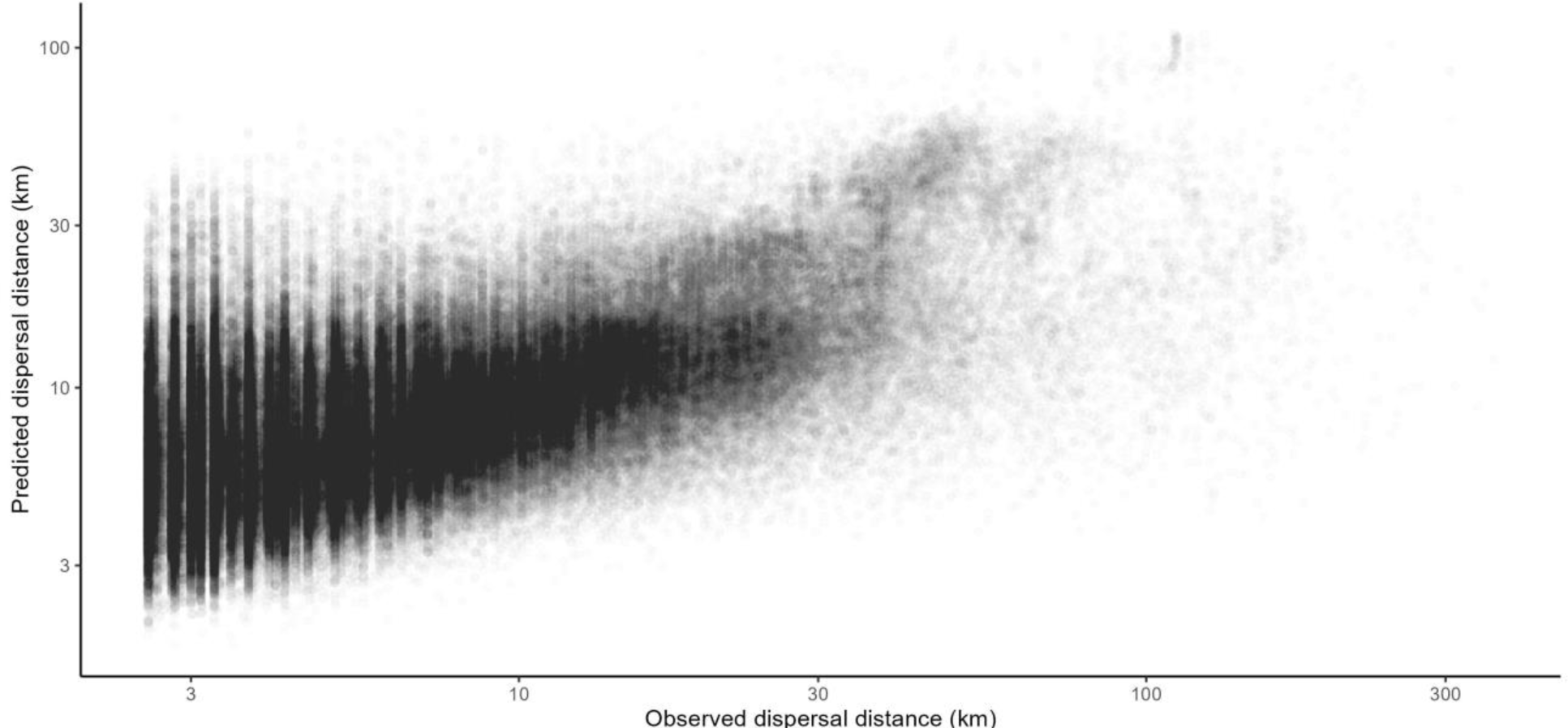
Predicted dispersal distances from the GLMM plotted against observed values.

**Table S1:**
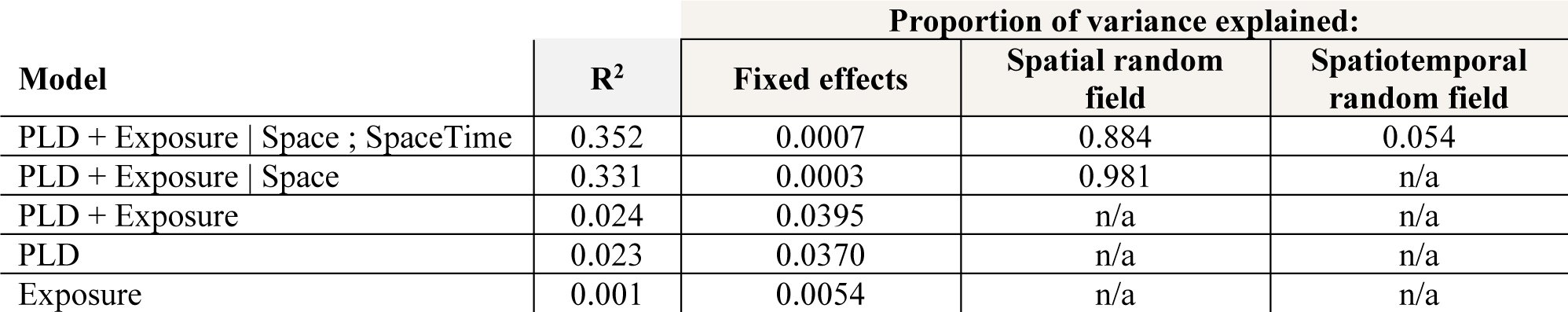
R^2^ and the proportion of variance explained by the fixed effects and random fields. In the model equations, Space indicates the spatial random field, and SpaceTime indicates the spatiotemporal random field. Proportion of variance explained (i.e. marginal and conditional R^2^) was calculated following methodology in Nakagawa and Schielzeth (2013). The random fields together explain 94% of the variance, and the fixed effects explain 0.07% of the variance. However, when the random fields are removed from the model, the fixed effects explain 4% of the variation. The inclusion of a spatial random field may absorb variation that would otherwise be explained by the fixed effects (Clayton et al. 1993).

